# Glycoinformatics approach for identifying target positions to inhibit initial binding of SARS-CoV-2 S1 protein to the host cell

**DOI:** 10.1101/2020.03.25.007898

**Authors:** Muhammet Uslupehlivan, Ecem Şener

## Abstract

COVID-19 outbreak is still threatening the public health. Therefore, in the middle of the pandemic, all kind of knowledge on SARS-CoV-2 may help us to find the solution. Determining the 3D structures of the proteins involved in host-pathogen interactions are of great importance in the fight against infection. Besides, post-translational modifications of the protein on 3D structure should be revealed in order to understand the protein function since these modifications are responsible for the host-pathogen interaction. Based on these, we predicted O-glycosylation and phosphorylation positions using full amino acid sequence of S1 protein. Candidate positions were further analyzed with enzyme binding activity, solvent accessibility, surface area parameters and the positions determined with high accuracy rate were used to design full 3D glycoprotein structure of the S1 protein using carbohydrate force field. In addition, the interaction between the C-type lectin CD209L and α-mannose residues was examined and carbohydrate recognition positions were predicted. We suggest these positions as a potential target for the inhibition of the initial binding of SARS-CoV-2 S1 protein to the host cell.

## Introduction

SARS-CoV-2 emerged in Wuhan, China in December 2019 and spread quickly across the world. It resulted in 575,444 confirmed cases and 26,654 deaths as to World Health Organization data as of 29 March 2020. Therefore, in the middle of the pandemic, all kind of knowledge on SARS-CoV-2 may help us to find the solution.

The coronavirus spike protein (S) plays a key role in the early steps of viral infection. S protein comprises of two sub-units S1 and S2 which are responsible for the binding of the host cell receptor and fusion of the cellular membrane, respectively [1,2,3]. S protein also contains furin cleavage site at the boundary between the S1-S2 sub-units which mediates the membrane fusion and virus infectivity [1,2]. It has been suggested that different domains within a single S protein could bind multiple alternative receptors. Although ACE2 (angiotensin-converting enzyme 2) is known to be the SARS-CoV receptor [2,4], CD209L, a C-type lectin that binds to high-mannose glycans on glycoproteins, has also been found to be as an alternative receptor for SARS-CoV [5]. However, recent studies have mainly focused on the S protein-ACE2 interaction, there is a paucity of information on the SARS-CoV-2 S1 protein-CD209L lectin interaction. It has been found that CD209L lectin binds to SARS-CoV, human coronavirus 229E, Ebola, Hepatitis C, HIV, Influenza virus glycoproteins and may mediate the endocytosis of pathogens [6-11]. Thus, identification of the carbohydrate-lectin interaction sites on the 3D structure for understanding of the initial binding of virus to a host cell is crucial. Besides, as post-translational modifications regulate the host-pathogen interaction [12], identifying SARS-CoV-2 S1 protein modifications may help us to inhibit initial binding of the virus. 3D crystal structure of SARS-CoV-2 S protein with N-glycosylation positions was identified but, sites such as furin cleavage and O-glycosylation are missing. In addition, the detailed structure of N-glycan units was not included in this model [13]. As glycan structures typically exist in solution or on proteins, it is a big challenge to characterize the 3D structure of glycoproteins experimentally. However, computational structural biology allows us to generate 3D protein and glycoprotein modelling with high accuracy rate using all amino acid sequence [14-16]. Herein, we predicted O-glycosylation and phosphorylation positions using full amino acid sequence of S1 protein. Candidate positions were further analyzed with enzyme binding activity, solvent accessibility, surface area parameters and the positions determined with high accuracy rate were used to design full 3D glycoprotein structure of the S1 protein using carbohydrate force field. In addition, the interaction between the C-type lectin and α-mannose residues was examined and carbohydrate recognition positions were predicted.

## Results and Discussion

Determining the 3D structures of the proteins involved in host-pathogen interactions are of great importance in the fight against infection. Besides, post-translational modifications of the protein on 3D structure should be revealed in order to understand the protein function since these modifications are responsible for the host-pathogen interaction. Therefore, researchers primarily have focused on the S protein structure of SARS-CoV-2 virus and revealed the 3D structure with N-glycosylation positions. But, the detailed structure of the N-glycan units in these sites and the O-glycosylation positions are not included in the current model [13]. Also, it is known that the initial binding of other coronaviruses to host cell occurs via the alternative receptor C-type lectin CD209L besides ACE2 protein [5]. Taken together, herein, we first demonstrated SARS-CoV-2 S1 protein high mannose N-glycan structure and its interaction with CD209L. When 3D glycoprotein structure of S1 protein analysed, N-glycosylation positions were found to be located mainly on the N-Terminal Domain (NTD) (Figure 1). Since the Receptor Binding Domain (RBD) is known to be involved in ACE2 binding, we suggest that N-glycan positions on NTD may associated with the binding of C-type lectin. C-type lectins interact with α-linked mannose residues on high mannose N-glycan structures [17]. There has been a structural study on C-type lectin-glycan interaction but terminal sugar is not α-mannose here [18]. Thus, in this study, the interaction between the C-type lectin and α-D-mannose was analysed and interface residues were identified. ZDOCK docking and PDBePISA interface interaction analysis showed in two models with high accuracy rate that mannose sugar interacts with C-type lectin at Met282, Lys307, and Ser345 positions with hydrogen bonding (Fig. 2). We suggest these positions as the α-D-mannose recognizing sites having function on the lectin-sugar binding.

**Figure 1.**
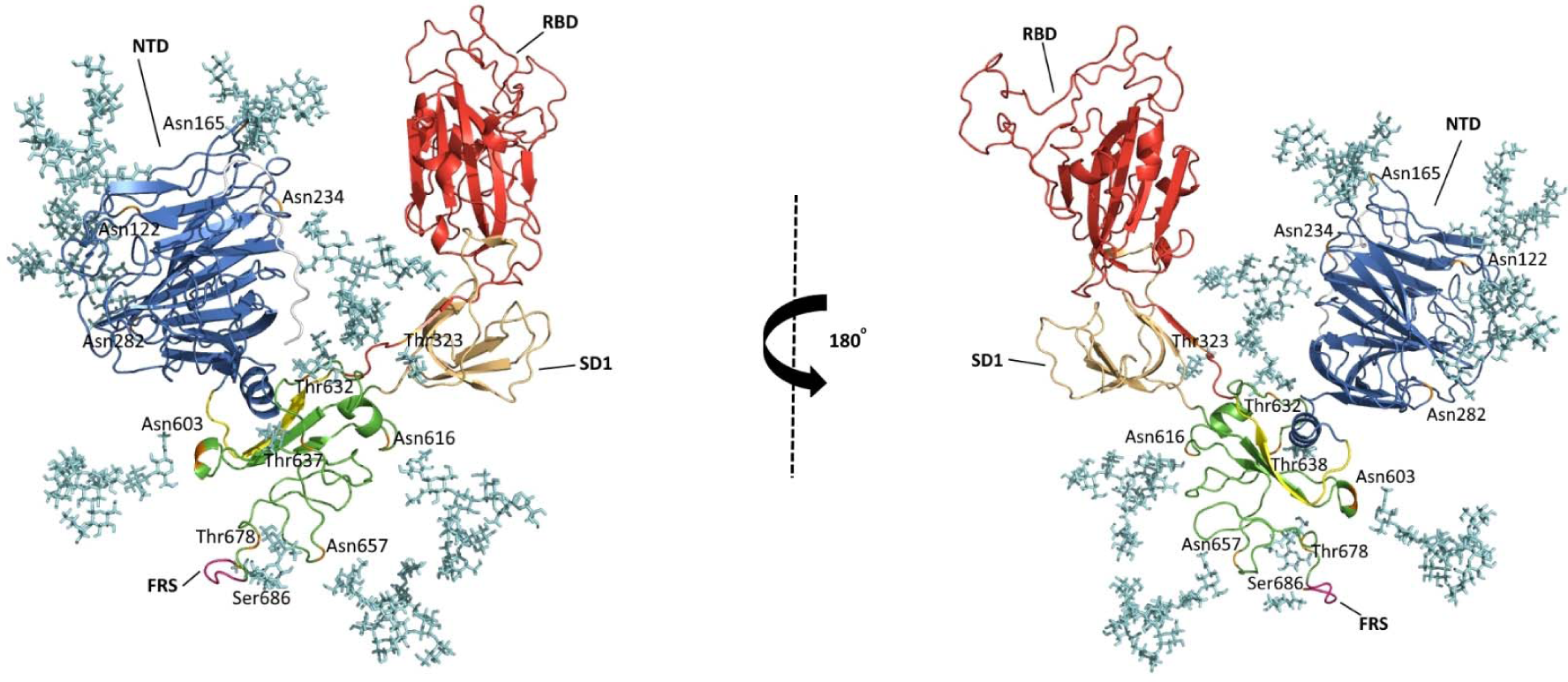
N- and O-glycosylated 3D glycoprotein structure of SARS-CoV-2 S1 protein. High mannose glycans of the N-glycosylation sites and core 1 type glycans of the O-GalNAcylation positions are shown in cyan color. N-glycosylation positions: Asn122, Asn165, Asn234, Asn282, Asn603, Asn616, Asn657; O-GalNAcylation positions: Thr632, Thr678; O-β-GlcNAcylation positions: Thr323, Thr638, Ser686. NTD: N-Terminal Domain, RBD: Receptor Binding Domain, FRS: Furin Recognizing Site. (The domain information was cited from Wong et al., 2004 and Wrapp et al., 2020.)

**Figure 2.**
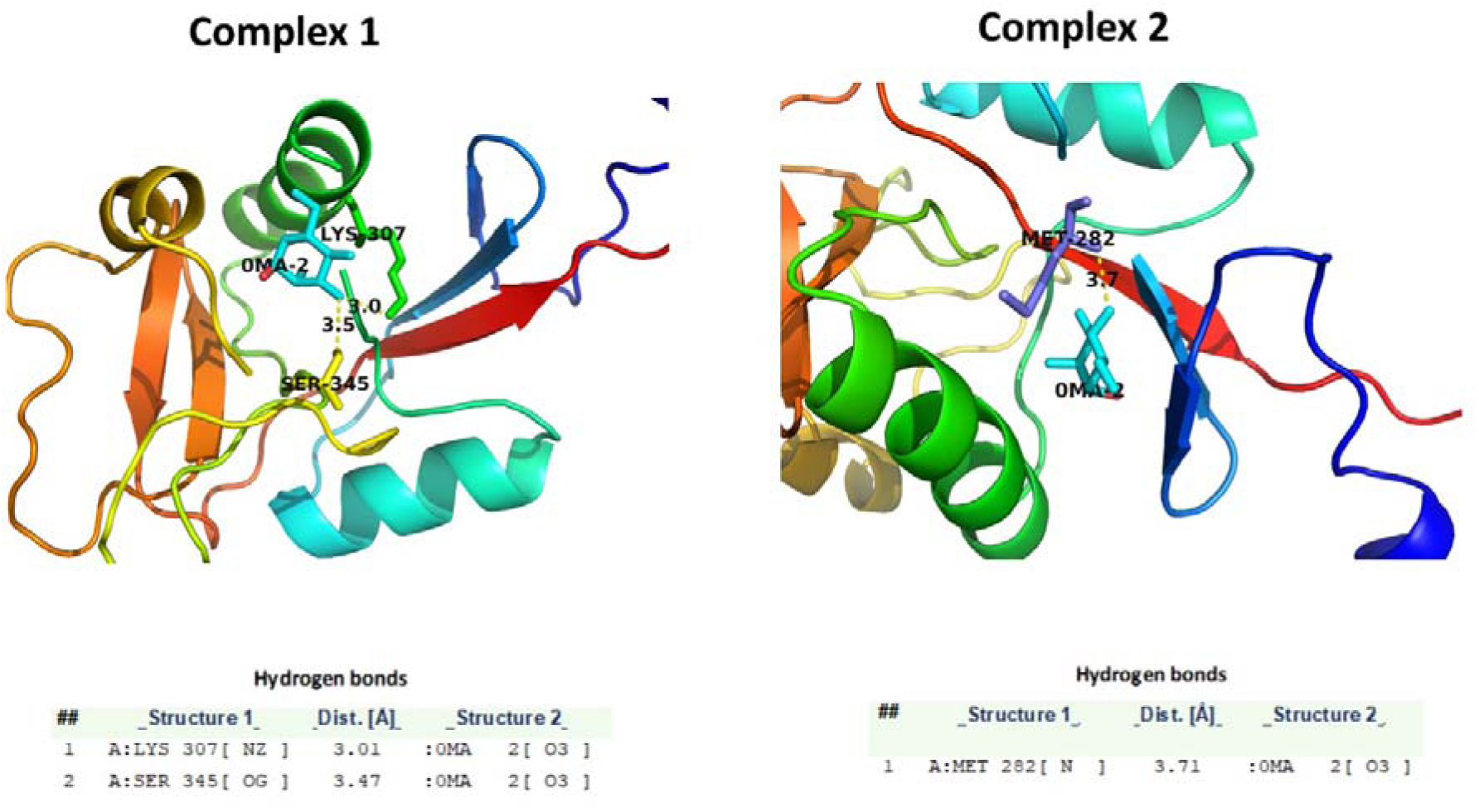
Receptor-ligand interactions of CD209L–α-D-mannose. ZDOCK and PDBePISA results showed that the 2 complexes have receptor-ligand interaction by hydrogen bonds at Lys307, Ser345, and Met282 positions. The mannose ligand is shown in cyan, receptor positions Lys307 in green, Ser345 in yellow and Met282 in blue color. All residues are shown in the stick representation.

Secondly, we identified O-GalNAc and O-GlcNAc modifications of SARS-CoV-2 S1 protein. O-glycans are known to be involved in protein stability and function [19]. The presence of O-glycans in some viral proteins has been demonstrated and suggested that glycans may play a role on biological activity of viral proteins. In the comparative study on human SARS-CoV-2 and other coronaviruses S proteins have showed that Ser673, Thr678, and Ser686 are conserved O-glycosylated positions and have suggested that SARS-CoV-2 S1 protein may show O-glycosylation at these positions [20]. In this study, S1 protein was found to be O-GalNAcylated at Thr632, Thr678 and O-β-GlcNAcylated at Thr323, Thr638, Ser686 positions (Table 1). When compared our results with the previous study, SARS-CoV-2 S1 protein found not to be O-glycosylated at Thr673, but O-GalNAcylated at Thr678 and O-β-GlcNAcylated at Ser686. Besides, we found additional O-glycosylation positions at Thr323, Thr632, and Thr638 on the S1 protein. On the 3D glycoprotein structure, Thr323 was found to be located at RBD and Thr678 and Ser686 located near Furin Recognizing Site (FRS) (Fig. 1 and 3). Since O-glycans are involved in protein-protein interactions [19], we suggest that O-glycans at Thr323 may play a role on binding to ACE2 and O-glycans at Thr678, Thr686 may responsible for furin protease enzyme binding. It has been known that O-glycans are responsible for the protein stability and creating mucin like domain as glycan shields involved in immunoevasion [21]. Based on this, we suggest that the O-glycosylation positions found may be involved in the stability of the S1 protein and immunoevasion of virus to the host cell.

**Table 1.**
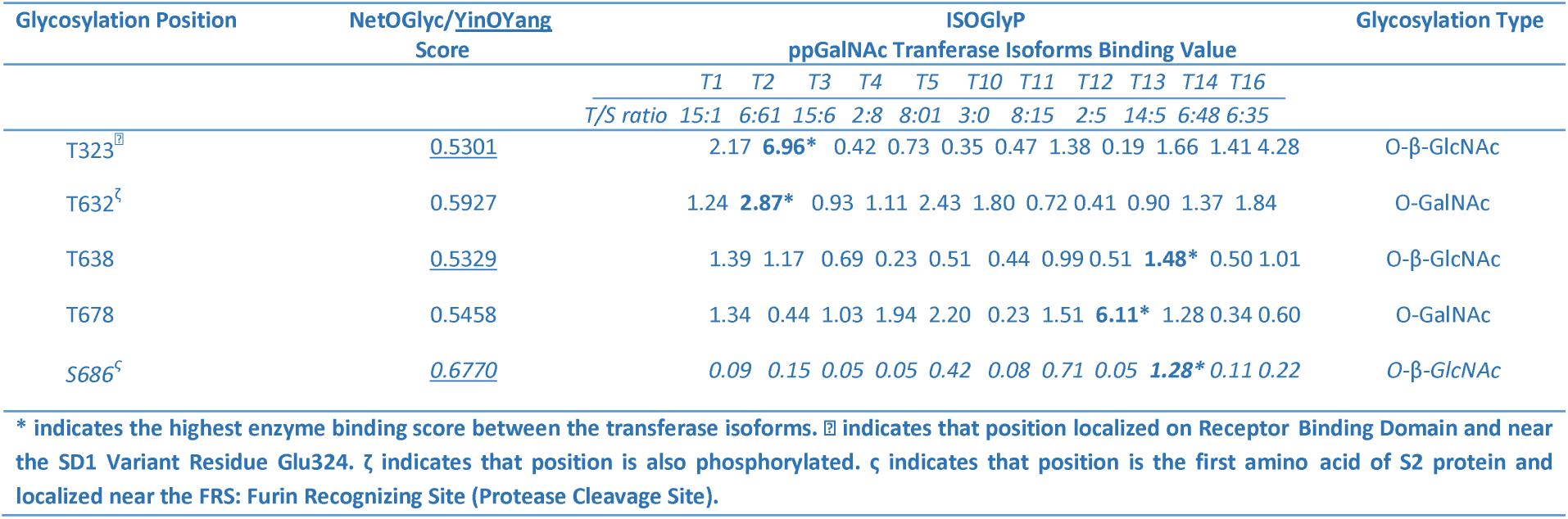
The predicted O-glycosylation positions of SARS-CoV-2 S1 protein.

**Figure 3.**
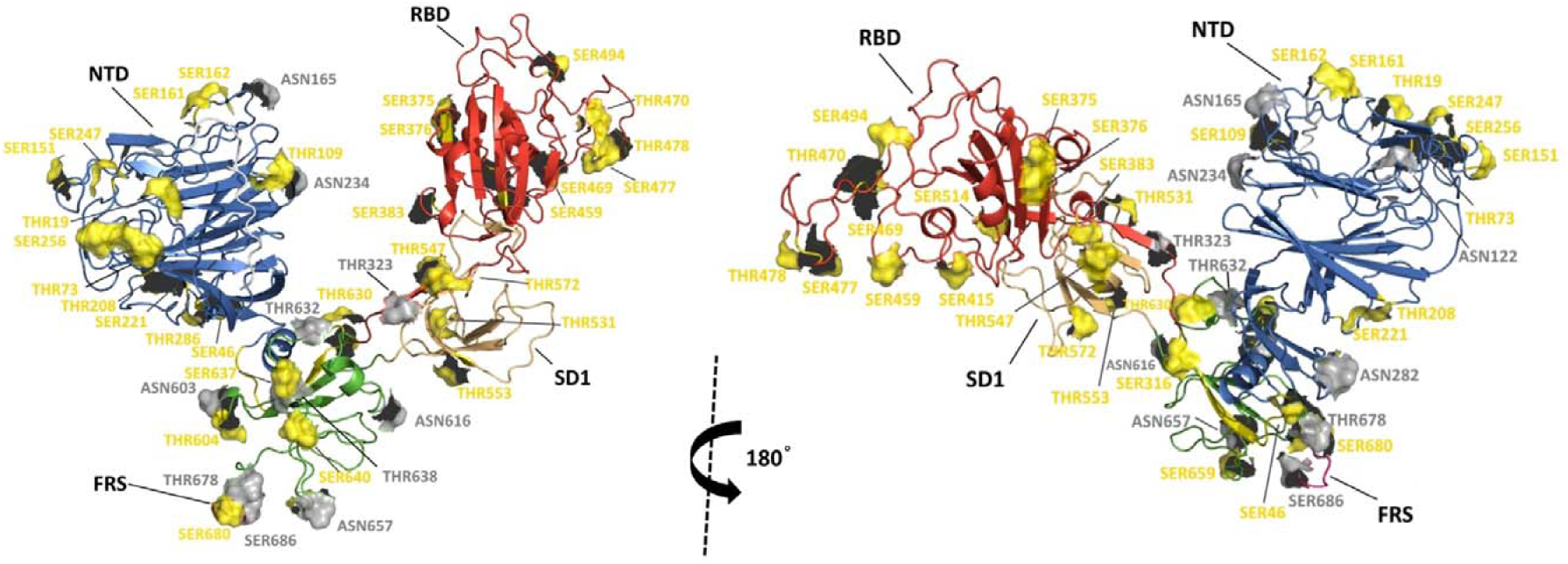
N-, O-glycosylation and Ser/Thr phosphorylation positions of SARS-CoV-2 S1 protein on the 3D structure. The glycosylation and phosphorylation positions are shown in grey and yellow, respectively. NTD: N-Terminal Domain, RBD: Receptor Binding Domain, FRS: Furin Recognizing Site

Phosphorylation of viral proteins are catalyzed by host cell enzymes like glycosylation modifications [12,22]. In this study, phosphorylation modification of SARS-CoV-2 S1 protein was also examined and 36 positions of Ser/Thr residues were found to be phosphorylated on almost all domains of the protein (Table 2). Davidson et al. have shown that S1 protein is phosphorylated at 7 positions [23]. Amongst them, Ser459, Ser637, Ser640 positions were found to be correlated with our results. Phosphorylation modification regulates the protein activity and function in cooperation with glycosylation [24,25]. When we examined the phosphorylation and glycosylation locations of S1 protein on the 3D structure; the phosphorylation positions at Ser161 and Ser162 were found to be located close to glycosylation position Asn165. Likewise, Thr109 with Asn234; Ser680 with Thr678 and FRS; Thr604 with Asn603; Thr630 with Thr632; Thr637 and Thr640 with Thr638; Ser659 with Asn657 were found to be located close to each other on the 3D structure (Fig. 3). Thus, we suggest these sites as critical sites for S1 protein activity.

**Table 2.**
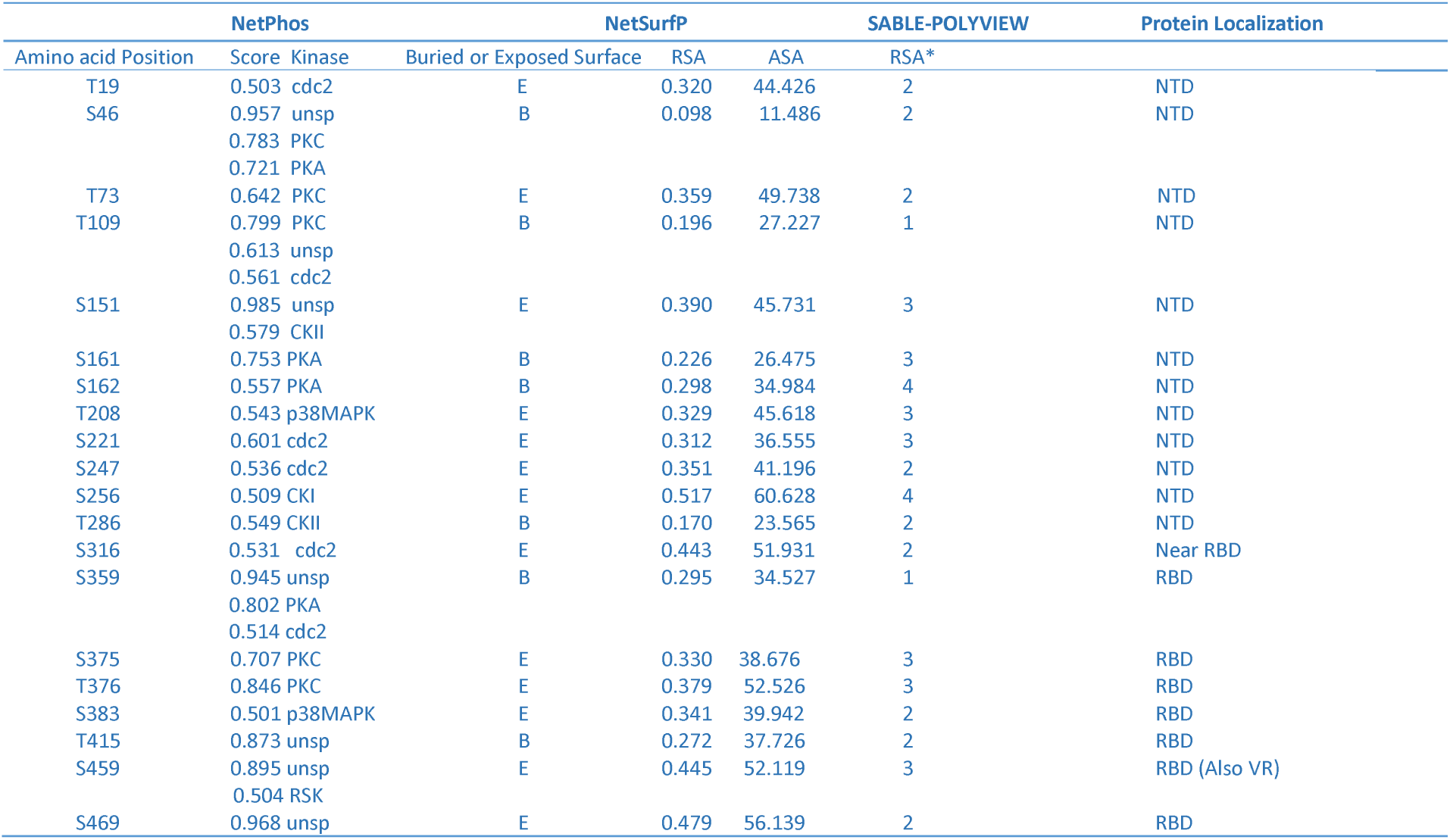
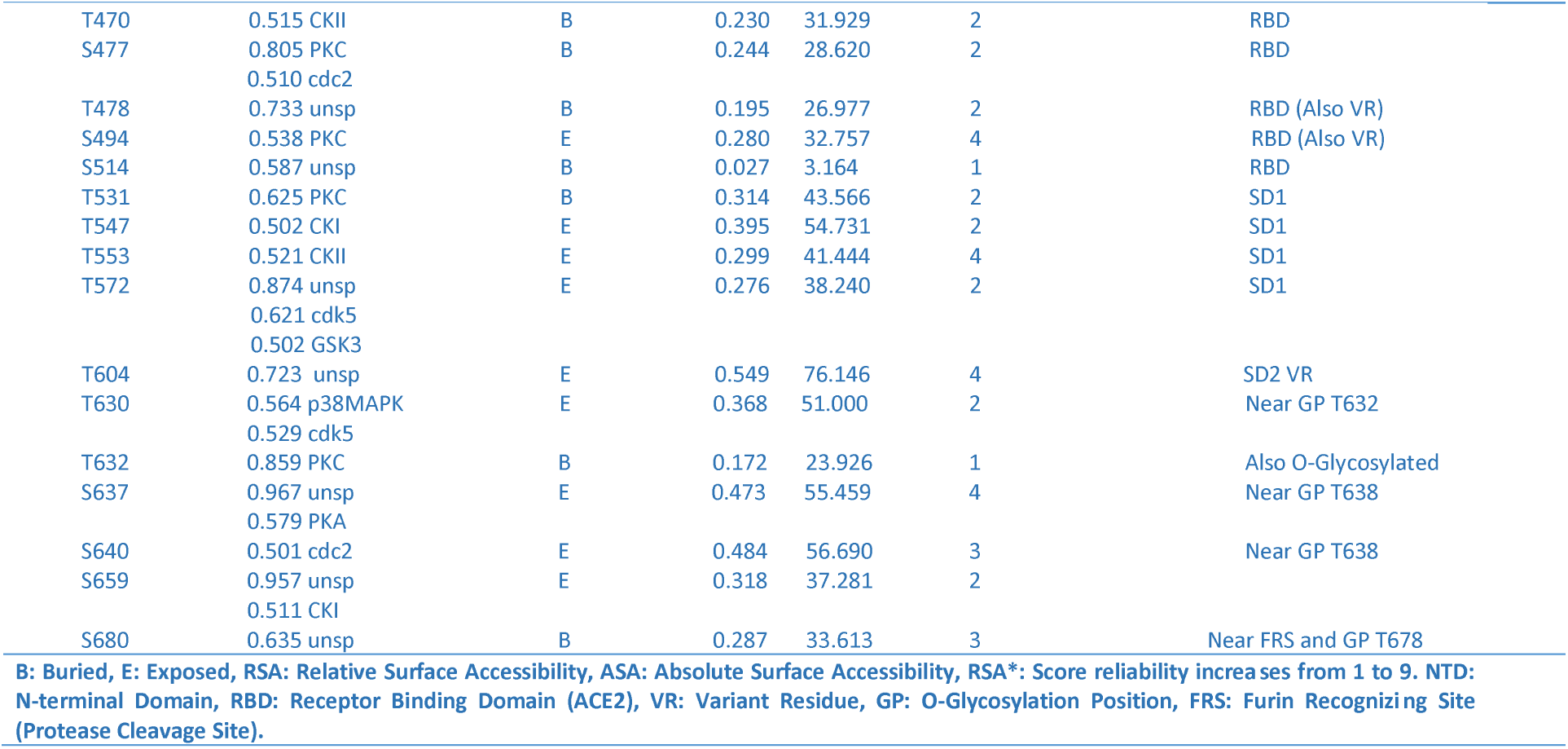
The predicted phosphorylation positions of SARS-CoV-2 S1 protein. All positions were analysed with the kinase binding, surface, and solvent accessibility parameters. The kinase specificity and protein localizations were also given in the table.

## Conclusion

COVID-19 outbreak is still threatening the public health. At this point, inhibiting the initial binding of virus to the host cell is crucial in order to find treatment. Since post-translational modifications regulate the host-pathogen interaction, identifying SARS-CoV-2 S1 protein modifications may help us to inhibit initial binding of the virus. Therefore, we focused on the CD209L lectin-α-D-mannose interaction of S1 protein and suggested Met282, Lys307, and Ser345 positions as targets for the initial binding. Also, we found that phosphorylation and glycosylation positions where located at close sites may be critical for S1 protein activity. Therefore, we suggest that, these positions can be targeted for drug development against COVID-19.

## Materials and Methods

### Prediction of O-Glycosylation and Phosphorylation Positions

Amino acid sequence of SARS-CoV-2 S1 protein was taken from NCBI with <u>QHD43416</u> ID. The potential O-glycosylation (O-GalNAc) and O-β-GlcNAc sites of S1 protein were analysed via NetOGlyc 4.0 [26] and YinOYang 1.2 Server [27], respectively. The threshold was chosen as 0.5 for both to predict high potential sites. The potential glycosylation positions which are found on NetOGlyc and YinOYang were analysed with the ISOGlyP server for the enzyme binding activity [28]. ISOGlyP was used to calculate all potential positions with ppGalNAc transferase isoforms which showed high enzyme binding activity and calculates the enhancement value product (EVP) values as an indication of glycosylation rates [29]. The potential phosphorylation positions of S1 protein was analysed with NetPhos 3.1 Server [30]. NetSurfP v1.1 was used to assess the surface and solvent accessibility of predicted Ser and Thr positions with glycosylation and phosphorylation positions [31]. The relative solvent accessibility of potential glycosylation positions was analysed using Poly-View 2D-SABLE protein structure prediction server [32].

### Glycoprotein Building on 3D Structure using Carbohydrate Force Field

3D structure models involving full amino acid sequence of S1 protein was taken with QHD43416 ID from I-TASSER (Iterative Threading ASSEmbly Refinement) [33]. The structure model has been generated by the C-I-TASSER pipeline, which utilizes deep convolutional neural-network based contact-map predictions to guide the I-TASSER fragment assembly simulations. GLYCAM-Web Server (AMBER carbohydrate force field) was used to screen highly reliable glycosylation sites on the protein 3D structure. GLYCAM-Web is dedicated to simplifying the prediction of three-dimensional structures of carbohydrates and macromolecular structures involving carbohydrates. Glycoprotein Builder runs SASA (Solvent Accessible Surface Area) prediction and finds the most appropriate glycosylation sites for adding glycan units on the 3D structure using molecular dynamics simulation [34,35]. Glycan units skeleton were chosen as Core 1 type that is the most common O-glycan motifs on proteins for O-glycan units. The common shape of high mannose type that is a conserved motif that plays an important role on the N-glycan formation were chosen for N-glycan units [17]. Additionally, all potential phosphorylation sites were analysed for SASA parameters on the 3D protein structure using GLYCAM and the positions with high accuracy rates were chosen as phosphorylation positions.

### Lectin-Carbohydrate Docking

In order to find out the CD209L lectin–mannose interaction, ZDOCK docking server was used [36]. CD209L lectin 3D structure (1.4 Å resolution) was taken from PDB (ID: 1XPH) [37] and α-D-mannose structure was taken from Glyco3D database [38]. PDBePISA server was used for the exploration of macromolecular interfaces between the receptor-ligand [39]. All structures were visualized with PyMOL.

